# Individual differences in frequency and topography of slow and fast sleep spindles

**DOI:** 10.1101/113373

**Authors:** Roy Cox, Anna C. Schapiro, Dara S. Manoach, Robert Stickgold

**Author notes:** Corresponding author: Roy Cox.

## Abstract

Sleep spindles are transient oscillatory waveforms occurring during non-rapid eye movement (NREM) sleep and are implicated in plasticity and memory processes. In humans, spindles can be classified as either slow or fast, but large individual differences in spindle frequency as well as methodological difficulties have hindered progress towards understanding their function. Using two nights of high-density electroencephalography recordings from healthy individuals, we first characterize the individual variability of NREM spectra and demonstrate the difficulty of determining subject-specific spindle frequencies. We then introduce a novel spatial filtering approach that can reliably separate subject-specific spindle activity into slow and fast components that are stable across nights and across N2 and N3 sleep. We then proceed to provide detailed analyses of the topographical expression of individualized slow and fast spindle activity. Group-level analyses conform to known spatial properties of spindles, but also uncover novel differences between sleep stages and spindle classes. Moreover, subject-specific examinations reveal that individual topographies show considerable variability that is stable across nights. Finally, we demonstrate that topographical maps depend nontrivially on the spindle metric employed. In sum, our findings indicate that group-level approaches mask substantial individual variability of spindle dynamics, in both the spectral and spatial domains. We suggest that leveraging, rather than ignoring, such differences may prove useful to further our understanding of the physiology and functional role of sleep spindles.

## Introduction

Sleep spindles are prominent rhythmic waveforms expressed by the mammalian brain during non-rapid eye movement (NREM) sleep. In humans, spindles are readily visible in the electroencephalogram (EEG) and are typically defined as short (0.5-2 s) bursts of activity in the sigma band (9–16 Hz). While they are initiated in the thalamus^1^, reciprocal interactions between cortex and thalamus shape their duration and amplitude^2,3^. Spindles are a defining feature of light N2 sleep, but also occur during deep N3 sleep where their occurrence is often obscured by large-amplitude ~1 Hz slow waves. Moreover, spindles can be present globally or restricted to specific brain regions^4–6^, propagate across the cortex^7^, and show complex patterns of interregional synchronization^8,9^. Functionally, sleep spindles are believed to be involved in processes of plasticity and offline memory consolidation^10–12^ as evidenced from relations between spindle activity and memory consolidation^13–16^.

Substantial evidence suggests that spindles can be classified as either slow (~10 Hz) or fast (~13 Hz). While it is presently unclear whether slow and fast spindles serve distinct functional roles^17–23^, they are associated with different hemodynamic sources^24^, respond differently to pharmacological interventions^25^, and preferentially occur in distinct phases of the slow wave^8,26–28^. Slow and fast spindles also have distinct EEG topographical distributions, with slow spindles having a more frontal expression and fast spindles occurring mostly centrally and parietally^29,30^. While these effects are well known at the group-level, little is known about the consistency of these topographical patterns across individuals. Given recent evidence of highly variable oscillatory profiles during wakefulness^31^, assessing whether similar variability exists during sleep is an important step towards a full understanding of the dynamics of sleep spindle activity within and across people.

Separating slow and fast spindles is nontrivial, in part because of the diversity of spectral definitions used by different groups for spindle detection. A non-exhaustive search of the literature reveals demarcation frequencies between slow and fast spindles placed at 12, 13, 13.5 and 14 Hz^17,24–26,32,33^. Similarly, the lower boundary of slow spindles has been set anywhere from 8 to 12 Hz^17,25,26,32,33^, and the higher boundary of fast spindles at 15 or 16 Hz^19,25^. Similar variability exists for studies not further differentiating between slow and fast spindles. This hinders comparison across studies.

The issue is exacerbated further by considerable variability in spindle frequency across individuals^34,35^. Subject-specific spindle frequencies are typically determined from peaks in the power spectrum, as such frequency domain peaks correspond to time-domain signals exceeding noise levels. However, slow and fast sigma peak frequencies are not confined to well-separated frequency ranges but form overlapping distributions at the group level^35^. Thus, even the demarcation line best separating slow and fast sigma peaks at the group-level likely results in incorrect classification of spindle activity for some subjects, and suboptimal separation for many others.

Individual differences in peak sigma frequencies and variable spectral spindle criteria together distort the correspondence between the oscillatory phenomena of interest and the measured activity used for subsequent analysis. This problem affects approaches investigating sigma power as a proxy for spindle activity^36^, as well as spindle detection algorithms based on band-pass filters and amplitude thresholds (e.g.,^18,37^).

To avoid these issues, approaches targeting subject-specific spindle frequencies have been developed^26,35,38,39^. However, while fast sigma peaks are typically prominent, slow sigma peaks are not always discernible, even at frontal channels where slow spindle activity is generally most pronounced^26^. This may reflect that slow and fast spindle topographies, while distinct, still show considerable overlap at both the level of sensors and underlying generators^27^. Moreover, as we will demonstrate, individual differences in spindle topography limit the effectiveness of selecting a single channel for slow sigma peak detection. Finally, slow spindles have been observed to express marked shifts of ~1 Hz between sleep stages, from faster in N2 to slower in N3^26^. In sum, available methods to separate spindle types do not always succeed and can lead to ambiguous results depending on what sleep stage is examined.

To overcome these difficulties, we introduce a novel approach to determine individualized slow and fast sigma frequencies in a data-driven fashion. The current report is organized into three main parts. In part one, we describe individual differences in the NREM power spectrum and the difficulties of detecting subject-specific sigma peaks from both channel-averaged and single-channel spectra. In part two, we detail a spatial filtering approach to facilitate isolation of subject-specific slow and fast sigma frequencies. We show that spectra derived from spatially filtered data allow for slow peak detection in more individuals and with less ambiguity than channel-based spectra. In part three, we characterize topographical aspects of slow and fast sleep spindle expression in N2 and N3, both to validate our sigma peak separation method, and to examine the commonalities, individual differences, and cross-night stability of spatially organized spindle activity. Part three will be particularly relevant to those interested in the implications of these methods for topographical spindle dynamics, and can be read separately from the first two parts. Combined, our results demonstrate the utility of our spindle separation approach and yield important new insights regarding the nature and variability of topographical sleep spindle dynamics across N2 and N3.

## Methods

### Protocol and participants

The current study utilizes two consecutive nights of full-night EEG data from 28 healthy individuals (age: 29.7 ± 6.0; 21 male, 7 female). These data were acquired as part of a double-blind, placebo-controlled, cross-over study of eszopiclone in schizophrenia patients^32,40^. Only the placebo nights are considered in the present report.

The study was approved by the Partners Human Research Committee. All participants gave written informed consent and were compensated monetarily for their participation. Participants had no personal history of mental illness as confirmed by screening with the SCID-Non-Patient Edition^41^. Furthermore, they reported no diagnosed sleep disorders, treatment with sleep medications, history of significant head injury or neurological illness, or history of substance abuse or dependence within the past six months. Upon completion of a pre-treatment visit to complete informed consent and undergo clinical and cognitive assessments, subjects received an actiwatch to wear from study enrollment to completion.

Subjects were randomly assigned to one of two treatment orders, placebo first or eszopiclone first, with a week in between the two treatment visits. Each of the two treatment visits consisted of two consecutive nights of polysomnography (PSG) monitoring at the Clinical Research Center of Massachusetts General Hospital. The first night of each visit served as a baseline night, while on the second night participants were trained for 12 min on a finger tapping Motor Sequence Task (MST)^42^ one hour prior to their usual bedtime. As no systematic differences in sleep architecture or spindle parameters were found between the baseline and learning nights, we focused our analyses on the first night and used to second night for validation and replication. On both nights of the placebo visit, participants received placebo at 10 PM. Lights were turned off at 10.30 PM and participants were allowed to sleep for up to 9.5 hours until they were woken up at 8 AM.

### Data acquisition and preprocessing

PSG was collected using 62-channel EEG caps (Easycap GmbH, Herrsching, Germany) with channel positions in accordance with the 10-20 system. Additionally, two single cup electrodes were placed on the mastoid processes, two around the eyes for electrooculography, two on the chin for electromyography, and a reference electrode was placed on the forehead. An AURA-LTM64 amplifier and TWin software were used for data acquisition (Grass Technologies). Impedances were kept below 25 kΩ and data were sampled at 400 Hz with hardware high-pass and low-pass filters at 0.1 and 133 Hz, respectively.

Sleep staging was performed in TWin using a limited number of channels with a contralateral mastoid reference on 30 s epochs according to standard criteria^43^. Initial processing of multi-channel EEG data was performed in BrainVision Analyzer2.0 (BrainProducts, Germany). All EEG channels were band-pass filtered between 0.3 and 35 Hz and notch filtered at 60 Hz. Channels displaying significant artifacts for more than 30 minutes of the recording were interpolated with spherical splines. EEG data were then re-referenced to the average of all EEG channels. Upon visual inspection, epochs containing artifacts were removed. To remove cardiac artifacts we used independent component analysis with the Infomax algorithm^44^. For each night and individual, remaining epochs were concatenated separately for the two sleep stages, resulting in 176 ± 61 (mean ± SD) and 211 ± 49 min of available N2 for the two nights (t(27)=-3,1, P=0.005), and 82 ± 44 and 85 ± 30 min of N3 (t(27)=-0.4, P=0.67).

All subsequent processing steps were performed in Matlab (the Mathworks, Natick, MA), using custom routines and several freely available toolboxes including EEGlab^45^ and Fieldtrip^46^. After removal of non-EEG channels and the mastoids, leaving 58 channels for analysis, we applied a surface Laplacian filter to each record^47^, also know as current source density estimation, as implemented in the CSD toolbox^48^. This approach served two purposes. First, the Laplacian renders data reference-free, thereby avoiding common interpretational issues related to the choice of reference. Second, this approach decreases the effects of volume conduction and accentuates local aspects of neural processing, thereby providing enhanced spatial precision for topographical analyses^49,50^. While the Laplacian is a spatial filter, we emphasize it is not the spatial filtering approach that we use to separate slow and fast sigma peaks.

### Power spectra and peak detection

After the Laplacian transformation, we determined the power spectrum for every epoch on every channel. In order to minimize the typical 1/f scaling of the spectrum, we obtained power estimates not from the Laplacian-transformed time series, but from its temporal derivative. This approach essentially multiplies power at every frequency bin by its frequency, thus counteracting the 1/f trend and allowing for easier detection of spectral peaks relative to surrounding frequencies^51^. Using the Laplacian-derivatives, we estimated power spectral density for each epoch using Welch’s method with 5 s windows and 50% overlap. We then normalized every electrode’s power spectrum during both N2 and N3 by dividing the spectrum by that electrode’s average power in the 0-4 Hz band across all N2 epochs. This normalization step was performed to enable direct comparison of power between sleep stages and individuals.

To determine global spectra, single-epoch spectra were averaged across all channels, before averaging across epochs, separately for N2 and N3. For visualization and peak detection, each individual’s 0-20 Hz spectra were rescaled between the minimum and maximum values in that range. Spectral peaks were detected using the Matlab *findpeaks* function with a minimum prominence setting of 0.01, where the prominence of a peak indicates how much a peak stands out as a function of both its intrinsic height and its location relative to other peaks. For our data, this setting corresponded to a very liberal detection threshold.

In addition to channel-averaged spectra, we determined spectra for two selected channels (frontal: Fz; parietal: Pz). We selected these channels based on evidence on the relative predominance of slow and fast spindles^26^. Single-channel spectra were averaged across epochs, again separately for N2 and N3 and both nights. Automated spectral peak detection was performed as before.

### Slow and fast sigma peak separation via spatial filters

In order to isolate each individual’s slow and fast sigma activity, we created linear spatial filters maximally enhancing slow vs. fast sigma activity, and vice versa. We then applied these filters to the multi-channel EEG time series to obtain a set of component time series that we analyzed in the frequency domain. In brief, the spatial filters were defined by eigenvectors extracted from covariance matrices, similar to principal component analysis (PCA)^52,53^. In a spatial filtering context, the PCA procedure operates on a single channel-by-channel covariance matrix and produces eigenvectors pointing in orthogonal directions that explain decreasing amounts of variance. This approach can be conceptualized as a "blind" source separation, in that resulting components are not necessarily physiologically meaningful. Rather than using a single covariance matrix as with PCA, we constructed two separate covariance matrices, one from slow sigma-filtered data and one from fast sigma-filtered data. We then performed generalized eigendecomposition (GED) to find eigenvectors maximally differentiating the two matrices. Contrasting with PCA, this technique may be viewed as a "guided" source separation procedure^52,53^ that spatially separates signal elements according to user-defined criteria. The GED approach has been used in various electrophysiological contexts, typically to maximize spectral power in one frequency band relative to broadband activity^52–55^. We here extend this notion by directly contrasting activity in two adjacent narrow-band ranges. Supplementary Matlab code demonstrates how to implement the GED analysis for several example sleep recordings.

In detail, we first band-pass filtered the Laplacian-transformed EEG separately in the slow (9-12 Hz) and fast (12-16 Hz) sigma ranges. We used Hamming-windowed finite impulse response filters (EEGlab: *pop_firrws*) with a high filter order (13,200) to create steep, narrow filters that have minimal overlap between the two passband ranges. After subtracting each filtered channel’s mean amplitude, we determined the "slow" covariance matrix **S** and the "fast" covariance matrix **F**, both of size 58 x 58 electrodes. If we designate **S** as the matrix whose signal we wish to accentuate and **F** as the matrix with the "noise", the eigendecomposition problem can be written as **SW** =**WFΛ**, where **W** is a matrix of eigenvectors and **Λ** is a diagonal matrix of eigenvalues^52,53^. In Matlab, **W** and **Λ** can be found via *[W,L] =eig(S,F).* The column in **W** with the highest corresponding eigenvalue then corresponds to the eigenvector that maximally enhances slow relative to fast sigma activity. Conversely, the eigenvector with the lowest eigenvalue has the opposite effect, maximizing fast relative to slow sigma power. Detailed treatments of the derivation of these equations can be found elsewhere^52–55^.

Although the most useful eigenvectors generally have relatively high and low eigenvalues, it is not known *a priori* which eigenvectors will yield the best results. We therefore multiplied the multi-channel, raw, broadband, Laplacian-transformed, EEG with the full matrix **W**, resulting in a time series of 58 components (where each component reflects a unique spatial weighting across all channels). These component time series, in turn, were transformed to the frequency domain. Similar to channel-based power spectra, we first took the temporal derivative of the multicomponent time series, and then estimated power spectral density using Welch’s method with 5 s windows and 50% overlap. We took the temporal derivative approach to reduce 1/f noise, and make channel-and component-based peak detection as similar as possible. However, we note that component peak location was not noticeably influenced by this procedure. Resulting component spectra were visualized and the first slow and the first fast component with peaks of sufficient quality were selected based on visual inspection. Typically, clear spectral sigma peaks were visible within the first ten (for slow sigma) or last ten (for fast sigma) components, with several components peaking at exactly the same frequency. Importantly, components were selected without knowledge of spectral peak frequencies in the corresponding channel-based spectra, guarding against experimenter bias. Finally, automated peak detection was performed as before to determine the precise frequency of spectral component peaks. This entire procedure was performed separately for each subject, for N2 and N3, and for both nights.

While spatial filters can be computed regardless of spectral structure, this technique does not create spectral peaks artificially. That is, in the absence of "true" band-limited peaks in the sigma range, spectra of returned components would not show clear peaks in the sigma range. Rather, such component spectra would have varied shapes that, as a result of the GED procedure, "happened" to be of overall larger amplitude in one of the two sigma ranges, but without a clear peak. Indeed, we observed such component spectra in a few subject/night/sleep-stage/spindle-class combinations, and we did not determine spectral peaks in these instances. We note that the spectral bands we used here for temporal filtering (9-12 and 12-16 Hz) are slightly different from the ones we later adopt as approximate slow (9-12.5 Hz) and fast (12.5-16 Hz) spindle ranges based on the distribution of sigma peak locations across individuals. However, we determined in several subjects that shifting initial filter ranges does not affect frequencies of subsequently identified component peaks by more than 0.2 Hz (also see *Supplementary Discussion*).

We defined an individual’s slow or fast sigma range as a 1.3 Hz window centered on his or her average sigma peak frequency across nights and sleep stages. This width was a compromise between a sufficiently broad range to capture small within-subject differences in peak frequency across sleep stages and nights, and sufficiently narrow to have non-overlapping slow and fast windows for as many individuals as possible. An additional consideration here was that for subsequent spindle detection (see *Spindle detection*), data needs to be filtered in each individual’s slow or fast sigma range. Thus, we also inspected each individual’s slow and fast sigma filters’ frequency response used for spindle detection. These inspections assisted both in arriving at the 1.3 Hz sigma ranges and in deciding which subjects to remove due to insufficient separation of the slow and fast ranges. Based on these considerations, we opted to exclude three subjects for subsequent analyses involving slow spindles, because their N2 and N3 slow sigma peaks differed by more than 0.7 Hz. Additionally, we excluded two subjects whose slow and fast sigma filters still overlapped. Of note, channel and component spectra for one of these subjects (S4) are shown in Fig. 2CG. Thus, these additional exclusions were not due to an inability to resolve closely spaced peaks with the GED component approach, but stemmed from an inability to adequately separate them with filter widths of 1.3 Hz.

### Topographical cluster analyses

For topographical analyses of sigma power, we averaged, per epoch and electrode, power across the 1.3 Hz frequency range corresponding to each subject’s individualized sigma peaks, before averaging across epochs. This was done separately for N2 and N3, and for each night. Permutation-based statistical analyses on topographical data (sigma power, as well as spindle density and spindle amplitude derived from automated spindle detection) were performed with Fieldtrip using cluster correction^56^. In several sets of analyses, we compared N2 and N3, and slow and fast spindle topographies for different spindle metrics. Using 1,000 iterations, the paired samples t statistic, a *clusteralpha* value of 0.1, and a significance threshold of 0.05, clusters were deemed significant at P < 0.025 for two-sided testing.

### Spectral and topographical similarity

We determined the within-subject similarity of each individual’s power spectra (e.g., between N2 and N3, between nights), and the topographical correspondence of each individual’s spindle expression (e.g., between slow and fast spindles, between N2 and N3) in conceptually and analytically similar ways. First, we calculated the Pearson correlation coefficient between a subject’s two relevant spectra, or two relevant topographies. For spectra, values were normalized power estimates at every frequency bin from 0-20 Hz (103 bins). For topographies, values were spindle activity estimates (e.g., sigma power) at each of the 58 electrodes. This yielded, across subjects and for each comparison, a set of *n* correlation coefficients and associated P values (where *n* is the number of individuals included in the particular analysis). To assess these results at the group-level, we performed a one-sample t test comparing the set of *n* correlation coefficients to zero. In addition, we adjusted the set of *n* P values for multiple comparisons using the False Discovery Rate^57^ and report the percentage of subjects showing above-chance similarity of spectral or spatial profiles.

Importantly, while within-subject correlation coefficients provide a useful index of "absolute" spectral or topographical similarity, they ignore how similar spectral or topographical profiles are across individuals. For example, average within-subject, cross-night correlation values of 0.8 do not indicate meaningful individual stability if average between-subject correlations are also 0.8. Conversely, within-subject correlations of only 0.4 signal substantial individual stability of spectra or topographies if between-subject correlations are only 0.2.

Second, therefore, we asked how well an individual’s spectral or topographical pattern would allow us to differentiate that subject from other subjects. To this end, we trained, for each comparison of interest, a k-nearest neighbor classifier^58^ on all subjects’ spectral/topographical patterns in one condition (e.g., N2), and tested it on unseen data from the other condition (e.g., N3). This is implemented in Matlab as *fitcknn.* We set *k* = 1, and used the correlation distance (1 - Pearson correlation) as the distance metric. In effect, this means each unseen test record is labeled as the one it is most highly correlated with from the training set. Classifier performance was calculated as the proportion of test cases that were assigned the correct label, and classifier significance was assessed using binomial tests. Thus, classifier performance provides an index of whether - and how much - spectral or topographical patterns are more stable within than across subjects, thereby complementing the within-subject similarity estimates afforded by Pearson correlation.

### Spindle detection

Individual sleep spindles were detected with an automated algorithm adapted from one we (RC) employed earlier^14,18^. For each subject and each channel, and separately for the slow and fast sigma ranges, the Laplacian-transformed signal was zero-phase band-pass filtered in that subject’s frequency range of interest. Specifically, we used Matlab’s *firls* function to design steep filters of order 3,000 with a 1.3 Hz passband and 0.5 Hz transition zones around the individualized center frequency. The sigma envelope was calculated as the magnitude of the Hilbert-transformed filtered signal, and was smoothed with a 200 ms moving average window. Whenever the envelope exceeded an upper threshold a potential spindle was detected. Crossings of a lower threshold before and after this point marked the beginning and end, respectively, of the spindle. Start and end points were required to be at least 400 ms and no more than 3,000 ms apart. Per channel, thresholds were set at the average N2 smoothed sigma amplitude envelope + 3 SDs (upper), and + 1 SD (lower). Spindle events were discarded whenever power in any 20-80 Hz frequency bin exceeded that of any frequency bin in that individual’s sigma range (suggesting a broadband power increase rather than band-specific spindle), or when the spindle’s average amplitude envelope was > 4 SD above the mean (indicative of an outlier). We calculated separate threshold settings for slow and fast spindles to optimally adapt to differences in slow and fast sigma amplitude. For both slow and fast spindles, the same N2-based thresholds were used across N2 and N3 to prevent confounding by different levels of sigma power across these sleep stages. We then determined spindle density (spindles per minute) and the mean peak amplitude of spindle events for each combination of channel, sleep stage, and spindle class.

## Results

### Channel-based analyses

#### Sigma peaks in channel-averaged spectra

To depict oscillatory activity across the cortex, we averaged 58-channel EEG spectra across all channels for every 30 s epoch, and then across epochs. Fig. 1A shows these channel-averaged spectra from night 1 for all 28 subjects during both N2 (magenta) and N3 (green) in the 0-20 Hz range. Fig. 1B shows each individual’s detected N2 and N3 peak locations as correspondingly colored dots with surface areas proportional to each peak’s prominence. In both panels A and B, thick dashed vertical lines placed at 9, 12.5 and 16 Hz indicate boundaries that best separate putative slow and fast sigma peaks across subjects. These boundaries are based on careful inspection of peak location distributions in the channel averaged-spectra, as well as the component-based spectra we present later, and agree well with those used by some other groups^26,35^.

**Figure 1.**
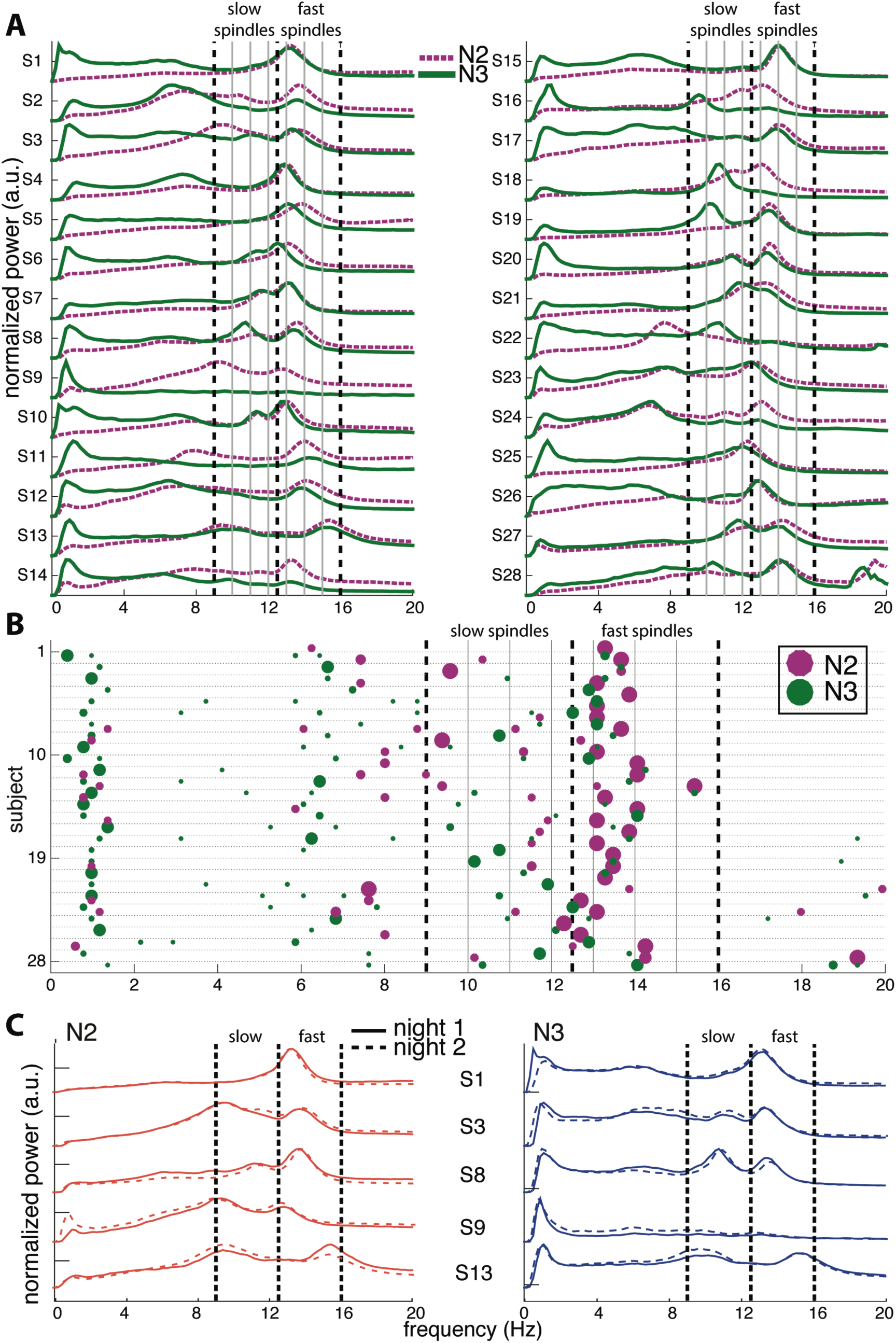
Variability of channel-averaged spectra and spectral peak locations in N2 and N3. (A) Individual subjects’ normalized spectra averaged across all channels are shown for N2 (magenta) and N3 (green) from night 1. For visualization, spectra have been rescaled to have the same amplitude range for every subject. Thick vertical dashed lines indicate slow and fast sigma boundaries at 9, 12.5 and 16 Hz. Thin solid vertical lines spaced at 1 Hz intervals assist in evaluating how alternative spectral definitions would partition sigma activity into slow and fast categories. (B) Spectral peak frequencies in N2 and N3 from night 1, corresponding to panel A. Each row represents an individual and colored dots indicate the location of spectral peaks for N2 (magenta, above line) and N3 (green, below line). Size of dots is proportional to peak prominence. Note the large variability in peak frequencies in the sigma range and the absence of clear slow sigma peaks for many individuals. In contrast, peaks in the 0.5-2 Hz range are highly consistent across individuals during N3. Vertical lines as in (A) (C) Normalized and rescaled channel-averaged N2 (left, orange) and N3 (right, blue) spectra of five example subjects from night 1 (solid) and night 2 (dashed). Individual differences in spectral shape are highly stable across nights. Vertical lines as in (A).

There was high variability in spectral profiles between subjects, especially in the prominence and number of visible peaks in the sigma range. Overall, we could identify one or more peaks in the broader 9-16 Hz range for all 28 subjects in both N2 and N3. In N2, we observed 13 individuals with just one peak, 14 with two, and one subject with three (S13), although there was typically one peak that was much more pronounced than the others. In N3, 12 individuals had just one identifiable peak, and 16 had two. Peaks in the fast sigma range (12.5-16 Hz) were relatively pronounced during both N2 (n=27) and N3 (n=24; including two subjects who had N3 peaks exactly at 12.5 Hz but corresponding N2 peaks > 12.5 Hz), as is evident from the relatively large dots in this range in Fig. 1B. In contrast, slow sigma peaks were often very shallow (small dots) or entirely absent in the channel-averaged spectra, resulting in identified peaks in far fewer subjects (N2: 16, including one subject with peak at 12.5 Hz but corresponding N3 peak < 12.5 Hz; N3: 20).

Importantly, there was considerable variation between subjects in the precise frequencies of slow (N2: 11.1 ± 1.0 Hz [mean ± SD]; N3: 10.9 ± 0.8 Hz; no stage difference: t(14)=1.2, P=0.24) and fast sigma peaks (N2: 13.5 ± 0.6 Hz; N3: 13.4 ± 0.6 Hz; trending to minimal stage difference: t(23)=1.9, P=0.07), without consistent separation into distinct slow and fast bands across subjects. We note that we set the sensitivity of peak detection very liberally to detect even small peaks, particularly in the slow sigma range. However, this enhanced sensitivity also resulted in detection of many shallow peaks in other frequency bands, especially in the adjacent theta range. In sum, these findings illustrate that two separate sigma peaks can be uncovered from channel-averaged spectra in roughly half the subjects.

N2 and N3 sigma profiles were often considerably different within the same individual (Fig. 1AB). This manifested in different ways across individuals and in different ways for slow and fast sigma activity. While relatively consistent for fast spindles, spectral peak frequencies did not correspond well between N2 and N3 in the slow sigma range (e.g., S3, S8, S18; note the disparity in location between pink N2 dots and green N3 dots). Indeed, for subjects having identifiable peaks in both stages, peak frequencies were highly correlated across N2 and N3 for fast sigma (R=0.91, P<10^-9^), but less robustly for slow sigma (R= 0.53, P=0.04). Thus, selection of slow and fast sigma bands has to address variability both across subjects and between sleep stages.

Of note, while the demarcation line at 12.5 Hz does a reasonable job for the group, some individuals’ peaks in Fig. 1A are centered right on, or very close to, that boundary (e.g., S4, S6, S23, and S25). We also note that many clear slow sigma peaks would have been missed entirely with lower slow sigma boundaries set at 11 Hz (e.g., S9, S16, S18, S19). Naturally, this issue is exacerbated as this threshold is moved further towards 12 Hz or beyond. Similarly, depending on precise boundary placement, closely spaced peaks could be lumped together into one spindle class. Thus, fixed frequency criteria have severe limitations

#### Stability and variability of channel-averaged spectra

Having observed a high degree of between-subject variability in NREM spectra, including sigma peak locations, we evaluated the stability of spectral profiles within individuals across night by comparing channel-averaged spectra from nights 1 and 2 (Fig. 1C; see Supplementary for spectra of all individuals). Spectral profiles were exceptionally similar across nights, including stable N2-N3 differences. To quantify this effect, we computed the degree of cross-night similarity as the Pearson correlation between night 1 and night 2 spectral profiles. We did this separately for N2 and N3. Across subjects, this yielded very high correlation coefficients of 0.97 ± 0.02 for N2 and 0.95 ± 0.05 for N3. These distributions were significantly different from zero (one-sample t-test: N2: t(27)=215, P<10^-44^; N3: t(27)=215, P<10^-35^), and spectra were more similar across nights for N2 than N3 (paired t-test: t(27)=2.8, P=0.009) Additionally, we adjusted individual subjects’ correlation P values for multiple comparisons using the False Discovery Rate^57^, and found that all individuals had P_corr_<0.05 for both N2 and N3.

While these findings demonstrate high within-subject similarity of spectral profiles, we also observed quite high correlations between spectra from different individuals (N2: 0.71 ± 0.16; N3: 0.57 ± 0.21), indicating NREM spectra show lower but substantial “baseline” levels of similarity between subjects. To emphasize the fact that there is individual spectral stability beyond between-subject levels, we asked if we could identify individuals based on the similarity of their power spectra across nights. To this end, we trained k-nearest neighbor classifiers on power spectra from one night and tested them on spectra from the other night, separately for N2 and N3. Given chance level performance of 3.6% (1/28), we obtained very high subject identification rates of 96% (27/28) for N2 (both directions: night 1->night 2 and night 2->night 1), and 82% (23/28) (night 1->night 2) and 89% (25/28) (night 2->night 1) for N3 (all P<10^-16^, binomial test). This robust stability of NREM sleep spectra underscores the trait-like nature of individual differences in oscillatory expression.

As noted above, we observed large differences between N2 and N3 spectral profiles within individuals (Fig. 1A). To quantify this effect, we computed the correlation between each individual’s N2 and N3 channel-averaged spectra. Across subjects, this yielded moderate correlation values (night 1: 0.45 ± 0.27; one-sample t-test: t(27)=8.9, P<10^-8^; 26/28 with P_corr_<0.05; night 2: 0.50 ± 0.21; one-sample t-test: t(27)=12.5, P<10^-11^; 24/28 with P_corr_<0.05). Again, however, these correlation values do not indicate whether spectra are more similar within than between individuals. To address this issue, we trained classifiers on N2 power spectra and tested them on N3 spectra from the same night, and vice versa. Resulting subject identification rates for classifiers trained on N2 and tested on N3 were modest at 18% (5/28, night 1) and 21% (6/28, night 2) (both P<10^-4^). Performance for classifiers trained on N3 and tested on N2 was even lower: 7% (2/28, P=0.08, night 1) and 11% (3/28, P=0.017, night 2). This suggests that, with a few exceptions, spectral profile shapes during N2 and N3 were generally as different within as between individuals.

In sum, while N2 and N3 spectra differ considerably within individuals, these differences are highly stable across nights. These data reinforce the notion that NREM spectra constitute robust individual traits^34,59^, and suggest the existence of similarly trait-like sigma peak frequencies. We return to this issue in more detail in the following sections.

#### Single-channel spectra

While averaging spectra across channels offers a topographically unbiased perspective, this approach may also obscure and distort spectral peaks present on a limited number of channels. Thus, the appearance of unitary sigma peaks in Fig. 1A for some subjects may reflect the merger of closely spaced slow and fast sigma peaks, or the attenuation of slow sigma peaks if they are only present on a few frontal channels.

We therefore turned to examining spectra on individual frontal (Fz) and parietal (Pz) channels where slow and fast spindles are most frequently reported^26^. Fig. 2A shows data for S10, with clearly separable slow frontal (blue) and fast parietal peaks (red), during both N2 (dashed) and N3 (solid). In a more ambiguous example (S6, Fig. 2B), posterior channel Pz exhibited clear peaks around 13 Hz, while frontal channel Fz showed a peak at 11.5 Hz during N3, presumably reflecting slow spindles. However, Fz displayed a much faster 12.9 Hz peak during N2, ostensibly signaling fast rather than slow spindle activity. In yet another case (S4, Fig. 2C), both the frontal and parietal channels showed peaks around 13 Hz with no suggestion whatsoever of a separate slow sigma peak. Thus, while examining channel-specific spectra may improve slow sigma peak detection in some cases, in many others it does not result in clearer peaks relative to the channel-averaged spectrum.

**Figure 2.**
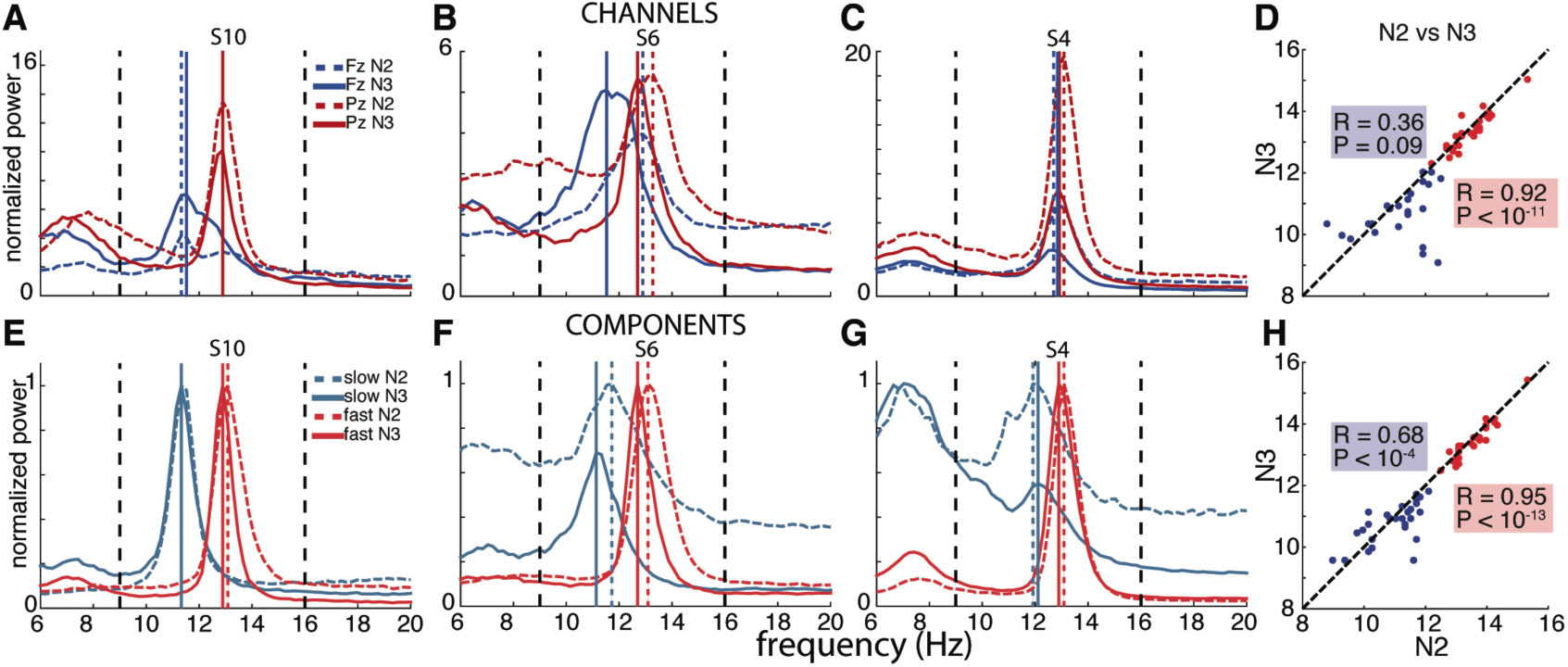
Channel-and component-based spectral peaks. (A-C) Single-channel spectra for three selected subjects for frontal channel Fz (blue) and posterior channel Pz (red), for N2 (dashed) and N3 (solid). Power obtained using derivative of Laplacian-transformed time series, and normalized to average 0-4 Hz power during N2. (D) Correspondence between channel-based peaks identified during N2 and N3, calculated separately for Fz and Pz. (E-G) Component-based spectra for same individuals as in panels A-C. Spectra scaled between 0 and 1 to account for varying amplitudes of different components. (H) Correspondence between component-based peaks identified during N2 and N3, calculated separately for slow and fast components. Correlation for slow sigma peaks is much greater compared to channel-based peak selection in D, indicating more stable estimates of underlying oscillatory frequency.

Nonetheless, we sought to identify peaks in the 9-12.5 Hz and 12.5-16 Hz bands on these channels where possible. Using the same peak detection algorithm as before, we identified fast peaks on Pz for all 28 subjects in both nights during N2, and 27 (first night) and 28 (second night) subjects during N3. Moreover, peak frequencies were highly correlated across nights, for both N2 (R=0.79, P<10^-6^) and N3 (R=0.92, P<10^-11^). In contrast, we could isolate peaks in the slow sigma range on Fz for only 19 and 22 subjects during N2, and 24 and 26 subjects during N3. Still, peak slow frequencies were significantly correlated across nights (N2: R=0.81, P<10^-4^; N3: R=0.79, P<10^-5^).

Given these high cross-night correlations for both slow and fast sigma, we averaged peak frequencies across nights, separately for N2 and N3. When a peak was only detected in one night, we used that value. This resulted, for slow sigma, in sample sizes of 23 for N2 and 26 for N3, while all 28 subjects could be included for fast sigma in both N2 and N3. Next, we examined the correspondence of peak frequencies between N2 and N3. As can be seen from Fig. 2D, this correlation was very high for fast sigma (R=0.92, P<10^-11^), with a small, but significant difference in frequency between N2 and N3, seen as points on average lying slightly below the identity line (13.4 ± 0.6 vs. 13.3 ± 0.6 Hz, t(27)=2.3, P=0.03). In contrast, for slow sigma we found no reliable association between stages (R=0.36, P=0.09), and observed a significant difference between N2 and N3 frequency that was much greater than for fast sigma (11.2 ± 1.0 vs. 10.7 ± 0.8 Hz; t(22)=2.2, P=0.04), differing in individual cases by up to 3.3 Hz. The latter result confirms previous observations that peak frequencies of slow sigma can differ substantially between N2 and N3^26^. These findings suggest that while individual fast sigma frequencies may be readily obtained from single-channel spectra, identifying a single slow sigma peak frequency for an individual is more problematic.

### Component-based analyses

Together, findings from the previous sections suggest that the identification of slow and fast spindle frequency bands either by the use of individual channels or by the use of all-channel averages is problematic. We sought to overcome these limitations by harnessing the spatiospectral structure inherent to multi-channel EEG recordings. Conceptually, the main obstacle to identifying slow sigma peaks is that, for any individual, it is not known *a priori* which channels express slow sigma most clearly and are least affected by interfering fast sigma activity. By determining an optimal combination of channels that enhances slow as opposed to fast sigma activity, and vice versa, the problem of identifying individual subjects’ slow and fast sigma frequencies becomes tractable.

Computationally, we derived the desired channel combinations using a linear spatial filtering approach based on generalized eigendecomposition (GED), a technique conceptually and mathematically related to principal component analysis. These spatial filters were then applied to the original EEG, resulting in "components" that we subsequently analyzed in the frequency domain (see Methods). Fig. 2EFG show the results of such analyses in three subjects. For S10, who already expressed clear slow and fast sigma peaks in single-channel spectra (Fig. 2A), the GED-based component approach identified highly similar peaks (Fig. 2E). More importantly, and in sharp contrast to the channel-based approach, clear slow peaks were also identified for both S6 and S4 during both N2 and N3 sleep (Fig. 2FG). For S6, where the frontal channel appeared sensitive to fast rather than slow spindle activity during N2 (Fig. 2B), components now identified slow peaks in both N2 and N3, with peaks frequencies only 0.6 Hz apart (Fig. 2F). Remarkably, whereas S4 showed no inkling of slow sigma activity based on channel Fz during either N2 or N3 (Fig. 2C), the decomposition approach identified components in the slow sigma range for both N2 and N3 with highly similar peak frequencies (Fig. 2G). Importantly, these slow components were distinguishable from components with peaks in the fast sigma range, suggesting this technique is able to identify closely spaced, but distinct, oscillatory rhythms.

Using this GED component method, we identified fast peaks for a total of 110 of 112 subject-night-sleep stage recordings (all but 2 second night N3 recordings), similar to channel-based detection. In contrast, for slow sigma, component-based peaks could now be isolated for all but one subject-night during N2 and three during N3, substantially increasing the number of subjects with respect to channel-based detection (by 42% and 22% for N2, and 17% and 0% for N3). As with channel-based peak detection, cross-night correlations were very high for fast sigma (N2: R=0.93, P<10^-11^; N3: R=0.94, P<10^-11^) and slow sigma peaks (N2: R=0.77, P<10^-5^; N3: R=0.59, P=0.0015), indicating this method identified components with similar spectral properties across nights.

Repeating the analysis performed on peaks identified from single channels, we averaged peak frequencies across nights (or, when a peak was detected on only one night, used that value, resulting in inclusion of all 28 subjects for both slow and fast sigma), and again examined the correspondence of sigma peaks between N2 and N3 (Fig. 2H). For fast sigma, peak frequencies were highly correlated as before (R=0.95, P<10^-13^), with a small, but significant difference between N2 and N3, similar to the channel-based approach (13.5 ± 0.6 vs. 13.4 ± 0.6 Hz, t(27)=2.7, P=0.01). For slow sigma, however, peaks now also showed a strong correlation between stages (R=0.68, P<10^-4^; cf. R=0.36, P=0.09 for single-channel analyses, above). Moreover, slow sigma frequency no longer differed significantly between N2 and N3 (11.0 ± 0.8 vs. 10.8 ± 0.7 Hz; t(27)=1,4, P=0.15). Thus, the GED component approach identified highly similar peaks in N2 and N3, suggesting that the same underlying oscillatory phenomena are present, and may be captured, during both light N2 and deep N3 sleep. We also directly compared sigma frequencies as determined from the single-channel and component approaches and found that peak frequencies are not substantially shifted by the component approach (Supplementary Text).

In total, unambiguous sigma peaks that were stable across N2 and N3 in both nights could be isolated for all 28 subjects for fast spindles, and for 25 individuals for slow spindles. Of note, applying the same strict criteria to channel-based spectra would result in a further loss of 9 subjects for slow spindle analyses, thus reducing by 75% the amount of data excluded from downstream slow spindle analyses. Mean fast spindle frequency was 13.5 ± 0.6 Hz, ranging from 12.5 to 15.4 Hz, while average slow spindle frequency was 10.9 ± 0.7, ranging from 9.3 to 12.0 Hz. These ranges are well in line with previous studies^26,35^. Not surprisingly, a paired t-test showed these frequencies to be significantly distinct (t(24)=12.2, P<10^-11^). Similar to previous reports^60^, we found no reliable correlation between slow and fast spindle frequencies (R=-0.30, P=0.15).

In sum, component-based sigma peak frequencies were highly replicable across nights and between sleep stages, for both fast, and crucially, for slow spindles. While single-channel spectra yielded similarly consistent results for fast sigma activity across nights and stages, these correspondences were much poorer for channel-based slow sigma activity. Together, these findings demonstrate that subject-specific spindle frequencies can be identified with higher accuracy using a GED-based spatial filtering approach. Supplementary Matlab code illustrates how to implement the GED-based sigma peak detection approach for several example sleep recordings.

### Topographical analyses

We now turn to topographical analyses of spindle activity defined by the individualized sigma peaks identified in the previous section. These examinations serve two main functions. First, given the novelty of the GED-based component approach to isolate subject-specific sigma frequencies, these analyses aim to provide crucial validation of our method. In order to demonstrate that the identified oscillatory frequencies correspond to physiological slow and fast spindles, individually targeted spindle activity should replicate key findings regarding slow frontal and fast centro-parietal spindle topography^29,30^. Second, taking advantage of the increased sensitivity afforded by our method, we take a detailed look at the spatial properties of sleep spindle activity, both at the group-level and by examining individual variability. By comparing slow and fast spindle topographies, during both N2 and N3, across nights, and finally, between several often-used metrics of spindle activity, we make several novel observations that expand our understanding of spindles.

#### Group-level slow and fast sigma power topographies

In order to carry out topographical analyses, we set individualized sigma ranges as a 1.3 Hz band centered on the mean peak frequency across N2 and N3. Due to overlapping slow and fast sigma ranges in two individuals, we included 26/28 individuals for fast spindle analyses and 24/28 for slow spindles (see Methods). We then averaged every electrode’s normalized power spectrum across frequency bins corresponding to that individual’s slow and fast sigma range. We performed this procedure separately for N2 and N3, and for each night. In what follows, we report data from night 1, unless otherwise stated.

Across subjects, fast sigma power showed a clear centro-parietal topography, both during N2 and N3 (Fig. 3AB). Moreover, performing topographical statistics with cluster correction, we found that fast sigma power was significantly elevated during N2 relative to N3 in one large cluster comprising all electrodes (P=0.002, Fig 3C), although the difference was largest at parietal sensors. In contrast, we saw distinctly different topographical profiles for slow sigma power. While during N3 slow spindles exhibited a clear frontal topography as expected (Fig. 3E), slow sigma activity during N2 was expressed in a bilateral fronto-central fashion, albeit with reduced power (Fig. 3D). Statistical analyses (Fig. 3F) revealed that slow sigma power was higher in N3 than N2 in a frontal cluster comprising 13 electrodes (P=0.07), whereas the reverse was found for a posterior cluster of 14 electrodes (P=0.04). The distinctly low slow sigma power at Fz during N2 is noteworthy, as this electrode is often used to quantify slow spindles^19,25^.

**Figure 3.**
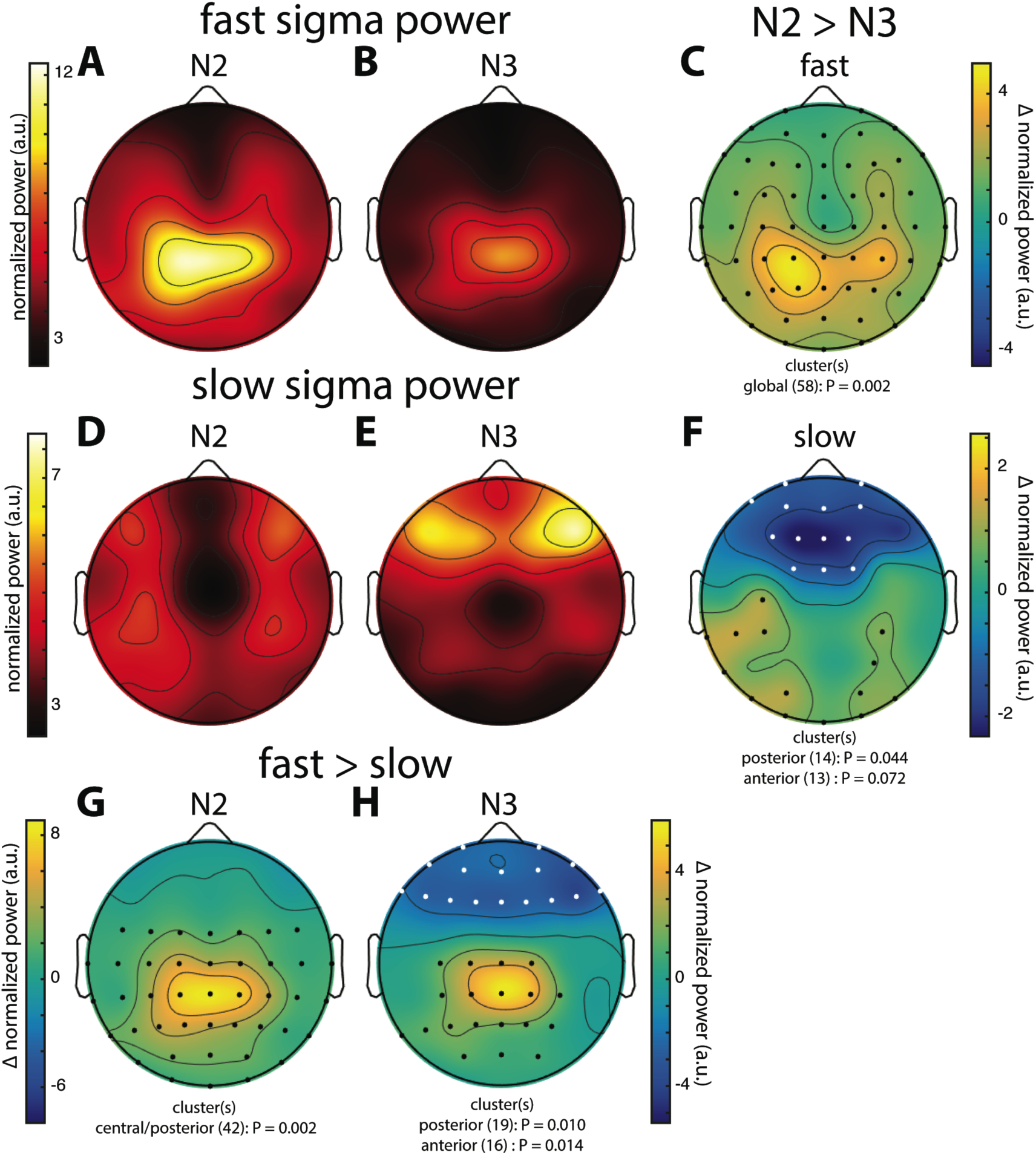
Group topographies of fast and slow sigma power during N2 and N3 from night 1. (A) N2 fast spindles. (B) N3 fast spindles. (C) N2 > N3 fast spindles difference. (D) N2 slow spindles. (E) N3 slow spindles. (F) N2 > N3 slow spindles difference. (G) N2 fast > slow difference. (H) N3 fast > slow difference. Significant electrodes from paired tests with cluster correction indicated on difference maps as black (positive) and white (negative) dots. Cluster size(s) (number of electrodes) and P value(s) indicated below each difference map. Note the different color scales for fast (AB) vs. slow spindles (DE), as well as different scales for each difference map.

We also directly contrasted fast and slow spindle activity and observed that, during N2, fast sigma power was significantly enhanced over a wide central-to-posterior region (P=0.002, Fig. 3G). In fact, while fast spindles clearly had a parietal focus, frontal fast spindle power during N2 was still numerically greater than slow sigma power. This observation could explain why N2 slow sigma peaks are not readily observed in channel data and may be easily overshadowed by fast sigma peaks. For N3, we similarly observed greater fast vs. slow spindle power over parietal regions (P=0.010, Fig. 3H), and, additionally, enhanced slow vs. fast power over a frontal cluster (P=0.014, Fig. 3H).

These group-level findings, based on our component analysis of individualized spindle frequency bands, confirm the differential topographical expression of frontal slow and parietal fast spindles, and N2 vs. N3 differences in fast spindle activity^26,30^. In addition, we report a more distributed slow spindle topography during N2 that was statistically distinct from the frontal pattern during N3. Thus, while slow spindles are most prominent over frontal areas during periods rich in slow waves, they extend to more parietal areas during light sleep. Importantly, we observed highly similar topographies and reproducible significant clusters when we analyzed data from night 2 (not shown). Encouraged that the individualized spatial filtering approach produced expected results, we decided to have a deeper look at the individual topographies contributing to the group effects.

#### Individual slow and fast sigma power topographies

Individual topographies of slow and fast sigma power from night 1 are shown for four subjects in Fig. 4, including two whose selected components were shown in Fig. 2. Inspection of scalp maps revealed both commonalities and considerable variability across subjects. We note that N2 and N3 topographies are plotted on the same scale to enable visual comparisons between sleep stages. In contrast, fast and slow spindle plots are on different scales to accommodate greater power in the fast sigma band, similar to the group-level observations.

**Figure 4.**
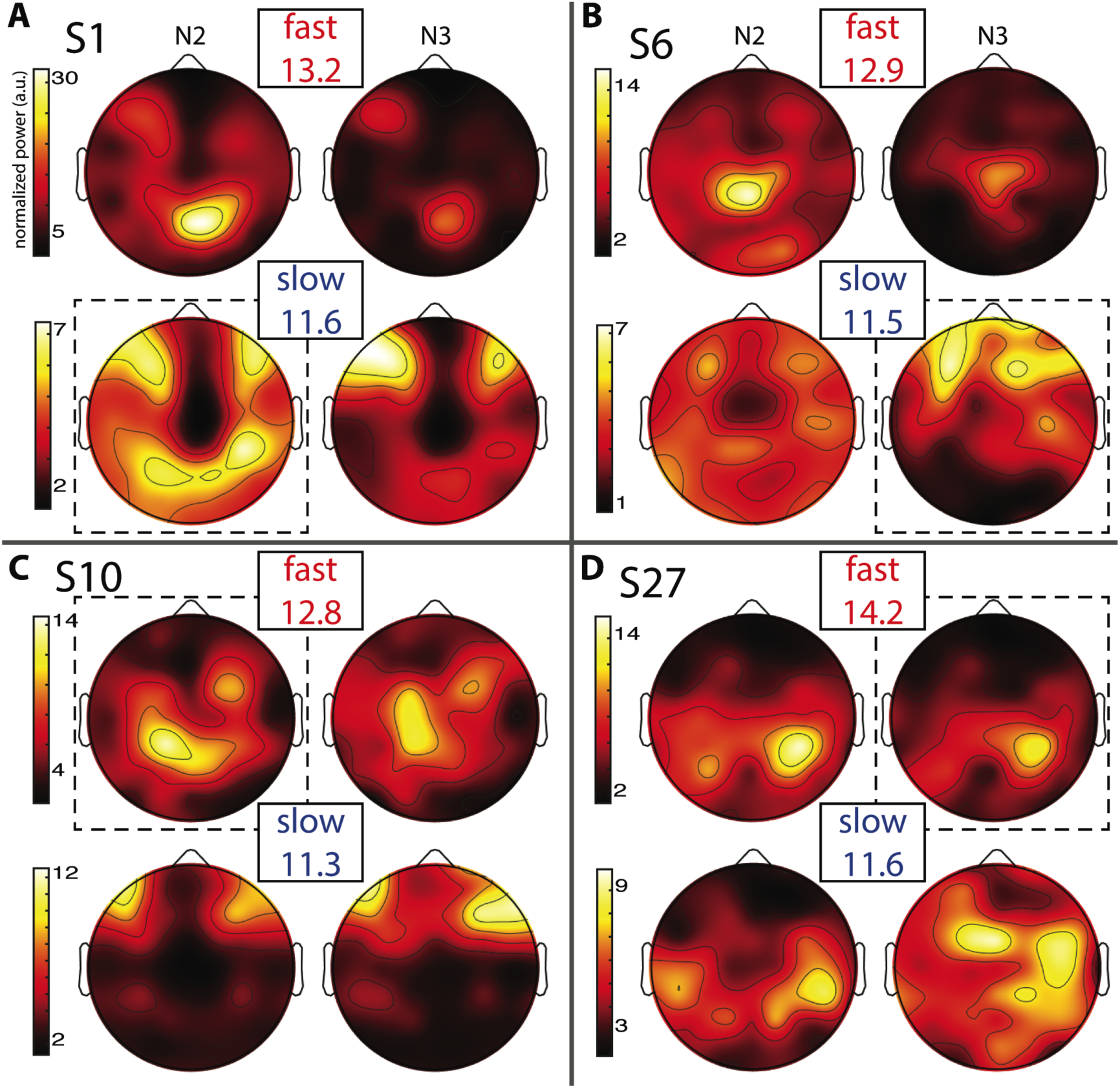
Individual topographies of fast and slow sigma power from night 1. For each subject panel, the top row displays normalized fast sigma power, the bottom row shows slow sigma power, and the left and right columns show N2 and N3 topographies, respectively. Numbers below “fast” and “slow” indicate each individual’s peak sigma frequencies in Hz. Dashed boxes indicate topographies used for cross-night comparisons of Fig. 5.

Fast spindles (top row in every panel) were expressed across the entire cortex, but with typical hotspots over central and parietal areas, although the exact topographies varied substantially. In line with group effects, fast sigma power was distinctly greater in N2 compared to N3 for some subjects (Fig. 4AB), but not others (Fig. 4CD). These findings also match the shape of individual power spectra in Fig. 1A. Regarding slow spindle activity, while three subjects exhibited a clear frontal topography during N3 (Fig. 4ABC), another had a more distributed, and distinctly non-frontal, pattern during deep sleep (Fig. 4D). During N2, slow sigma topographies were even more variable, with one subject having a frontal distribution matching the one seen in N3 (Fig. 4C), a second having both frontal and parietal hotspots (Fig. 4A), another having just posterior clusters (Fig. 4D), and yet another having a rather distributed profile (Fig. 4B). We note that the observed individual topographical variability emphasizes the difficulty of detecting slow sigma peaks from a single frontal channel. For example, S1 (Fig. 4A) did not express much slow spindle power at channel Fz, and, additionally, had higher fast than slow sigma power even over frontal regions (note the different color scales for slow and fast sigma). Indeed, this subject only demonstrated one unified sigma peak in the power spectrum of Fig. 1.

Despite overall power differences, each individual’s N2 and N3 topographies appeared quite similar within the same slow or fast spindle class. In order to quantify this, we correlated N2 and N3 sigma power across all electrodes for each individual, separately for slow and fast spindle classes. Thus, this approach assesses the similarity between topographical N2 and N3 patterns within subjects, irrespective of potential stage-dependent differences in absolute power. We found quite strong correlations of around 0.5 and 0.8 for slow and fast sigma, respectively (Table 1, *N2 vs. N3).* These values were significantly greater than zero both at the group level and for at least 75% of individuals across N2 and N3. Analogous to our approach for spectral stability, however, these results do not indicate whether similar correspondences might be found when using N2 and N3 sigma profiles from different subjects. We therefore turned to classifiers and asked if knowing individuals’ N2 sigma profiles would allow us to recognize them from their N3 sigma topography, and vice versa. Recognition rates were highly significant, at approximately 45% and 75% for slow and fast sigma (Table 1, *N2 vs. N3*). Thus, these findings suggest the same underlying subject-specific spindle generators are active across light N2 and deep N3 sleep states, most clearly for fast spindles.

**Table 1.**
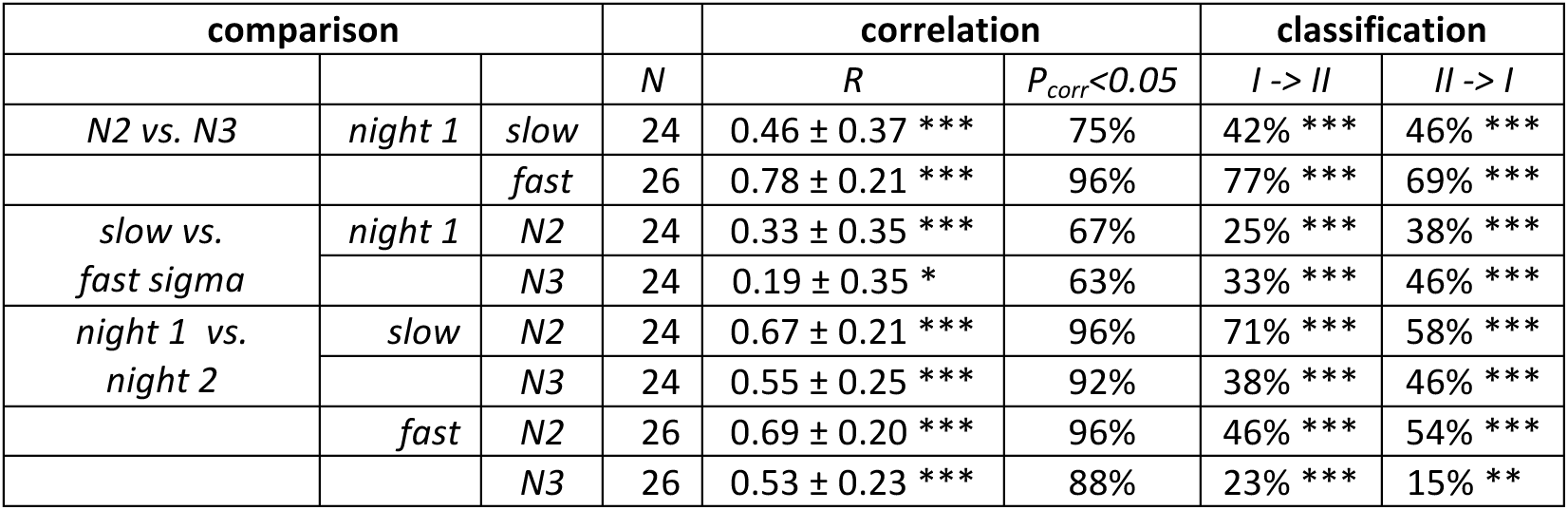
Within-subject similarity of sigma topographies between sleep stages, between spindle classes, and between nights. R: Pearson correlation coefficient (mean ± SD, across individuals); P_corr_<0.05: percentage of subjects with False Discovery Rate-corrected P values < 0.05; I-> II and II->I: classification direction. Significance levels (correlation: one-sample t-test vs. zero; classification: binomial test) indicated by *:< 0.05; **:< 0.01; ***:< 0.001.

As evident from Fig. 4, each individual’s slow and fast sigma topographies appeared to correspond poorly. Quantifying the degree of similarity between these spatial profiles, we found that slow and fast sigma topographies had average correlation coefficients of 0.3 and 0.2 during N2 and N3, respectively, indicating only limited within-subject correspondence of slow and fast sigma expression across the scalp (Table 1, *slow vs. fast sigma*). Again, however, these results do not account for the baseline similarity of slow and fast sigma profiles from different subjects. Using classifiers, we asked if knowing individuals’ slow sigma profiles would allow us to recognize them from their fast sigma topography, and vice versa. Across N2 and N3, we obtained significantly above chance cross-spindle-type classification rates of around 30% when trained on slow sigma and tested on fast sigma, and of around 40% when trained on fast sigma and tested on slow (Table 1, *slow vs. fast sigma*). These findings indicate that, even though an individual’s spatial profiles of slow and fast sigma expression are quite distinct (low correlation), they still share a sufficient degree of commonality to allow differentiation from other individuals’ sigma topographies in at least a third of our subjects. Overall, however, individual slow and fast sigma profiles were much less similar than N2 and N3 topographies of the same spindle type, suggesting these spindle classes are largely distinct.

So far, it is unclear whether individual differences in topographical expression of sigma power constitute stable traits, or rather, reflect night-specific differences in brain state that may be equally pronounced for the same individual across nights. As shown for the examples in Fig. 5, however, individual spatial differences were highly stable across nights, for both slow and fast sigma power, and for both N2 and N3. To quantify this effect, we next correlated sigma power across all electrodes between the two nights for each individual. We found substantial evidence for individual topographical stability across nights, with correlation values of around 0.6 for both slow and fast sigma topographies, and during both N2 and N3 (Table 1, *night 1 vs. night 2*). Note how these cross-night correlations were much higher than cros-sspindle-type correlations within the same night. We again turned to classifiers to ask if individuals can be recognized across nights based on their spatial expression of sigma activity. Recognition performance was significantly above chance for all analyses, but successful up to three times as many individuals for N2 than N3, and higher for slow than fast sigma profiles (Table 1, *night 1 vs. night2*). Whereas cross-night N3 fast sigma topographies were only slightly more similar within than between individuals, leading to classification rates around 20%, this difference was much greater for N2 slow sigma profiles, yielding recognition accuracy around 65%. The latter observation is noteworthy given that slow sigma peaks were most difficult to isolate from channel-based spectra in this sleep stage, indicating that meaningful slow sigma topographies may be extracted even under taxing circumstances.

**Figure 5.**
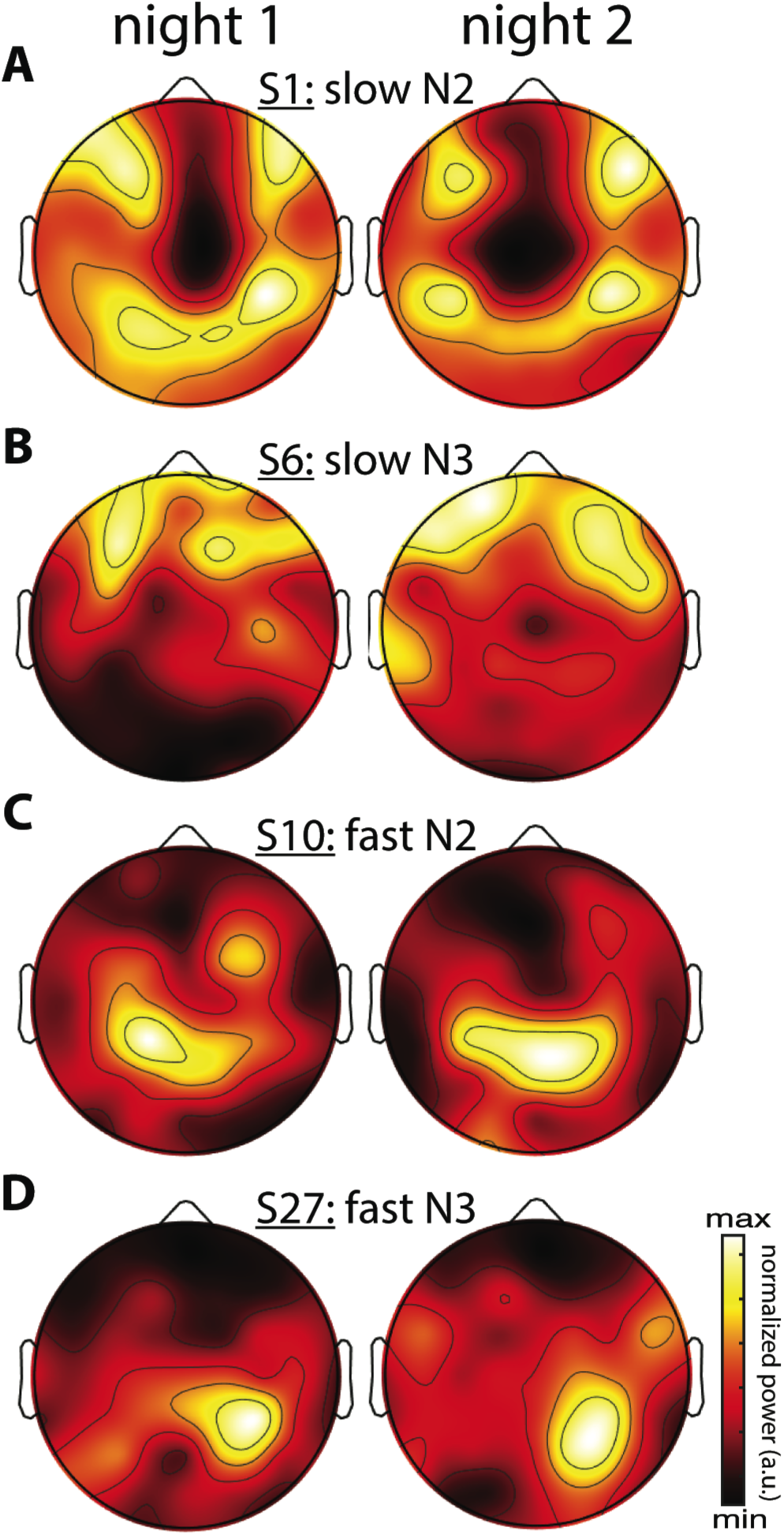
Stability of individual differences in sigma power topographies across nights. (A)Slow N2 sigma for subject S1. (B) Slow N3 sigma for subject S6. (C) Fast N2 sigma for subject S10. (D) Fast N3 sigma for subject S27. To emphasize topographical similarity, color scales were scaled between minimum and maximum value for each topography.

For all classification analyses reported in Table 1, it should be mentioned that recognition scores are influenced by both individual stability and between-subject variability. Higher “baseline” topographical similarity between different individuals for a particular sleep stage or spindle class could make it more difficult to differentiate between individuals, even if within-subject topographies are highly similar. Thus, the classifier approach takes between-subject variability into account, complementing the topographical correlation analyses that do not.

Together, these subject-specific analyses demonstrate that individual patterns of sigma expression are relatively stable across sleep states, but much less so across slow and fast sigma bands, providing additional support for the existence of two distinct spindle classes. Moreover, individual variability in topographical sigma expression is stable across nights, most prominently for N2 sleep. As such, individual topographical variability of both slow and fast spindles appears to reflect another individual trait of NREM sleep, similar to the cross-night stability of power spectra. In sum, while the previous section demonstrated clear indications of canonical slow and fast sigma topographies across subjects, these group effects mask substantial individual variability that must be taken into consideration when analyzing the spatial properties of spindle activity.

#### Topographies of detected spindles across and within subjects

A potential concern from the previous analyses is that by focusing on sigma power we do not treat spindles as distinct events with designated starts and ends. More generally, it is often assumed, at least implicitly, that sigma power and properties of individually detected spindles (e.g., spindle density, spindle amplitude) capture largely similar aspects of underlying spindle activity. However, as we are unaware of studies directly comparing these metrics, we next turn to examinations of group-level and subject-specific topographies of discrete spindle properties, and relate these to the sigma power patterns described in the previous sections.

We used an automated spindle detector^18^ to isolate individual sleep spindles using the subject-specific frequencies identified earlier. Briefly, the algorithm filters Laplacian-transformed channel data in the individualized sigma range of interest, and applies upper and lower thresholds based on the mean and standard deviation of the N2 sigma envelope (see Methods). Of note, this threshold approach dynamically adapts to the level of sigma signal present at each channel, thus contrasting with the sigma power approach that expressly does not account for such differences. Moreover, by running the algorithm twice, targeting slow and fast sigma ranges separately, different thresholds are applied for slow and fast spindle detection, aiding in isolating different-amplitude spindles. For example, upper thresholds were significantly higher across subjects for slow vs. fast spindle detection on channel Fz (0.20 ± 0.06 vs. 0.18 ± 0.07 μν/cm^2^, t(27)=3.6, P=0.001), but this relation was reversed on channel Pz (0.17 ± 0.07 vs. 0.21 ± 0.10 μV/cm^2^; t(27)=-2.4, P=0.02). After detecting spindle events in this manner, we calculated spindle density (number per minute) and mean spindle peak amplitude for each channel and individual, separately for N2 and N3, and separately for slow and fast spindles.

In general, group topographies of spindle density and peak spindle amplitude were consistent with sigma power profiles presented in Fig. 3. Compared to sigma power, spindle density profiles generally had a more diffuse appearance (Fig. 6ABDE), but topographical statistics indicated N2 vs. N3 differences for both fast (Fig. 6C) and slow spindles (Fig. 6F) similar to those seen for sigma power. Similarly, slow vs. fast spindle density topographies matched those observed for sigma power (Fig. 6 GH; noting that different thresholds were applied for detecting these two spindle types). Spindle amplitude topographies also looked similar to sigma power patterns, albeit with a more focal appearance (Fig. 6 IJLM). Similar to the other metrics, fast spindle amplitude was greater in N2 than N3 (Fig. 6K). In addition, slow spindles were also of significantly higher amplitude in N2 vs. N3 (Fig. 6N), including frontal regions where values were substantially lower in N2 for sigma power (significantly) and spindle density (numerically). Finally, fast and slow spindle amplitude topographies differed significantly as the other metrics did (Fig. 6OP).

**Figure 6.**
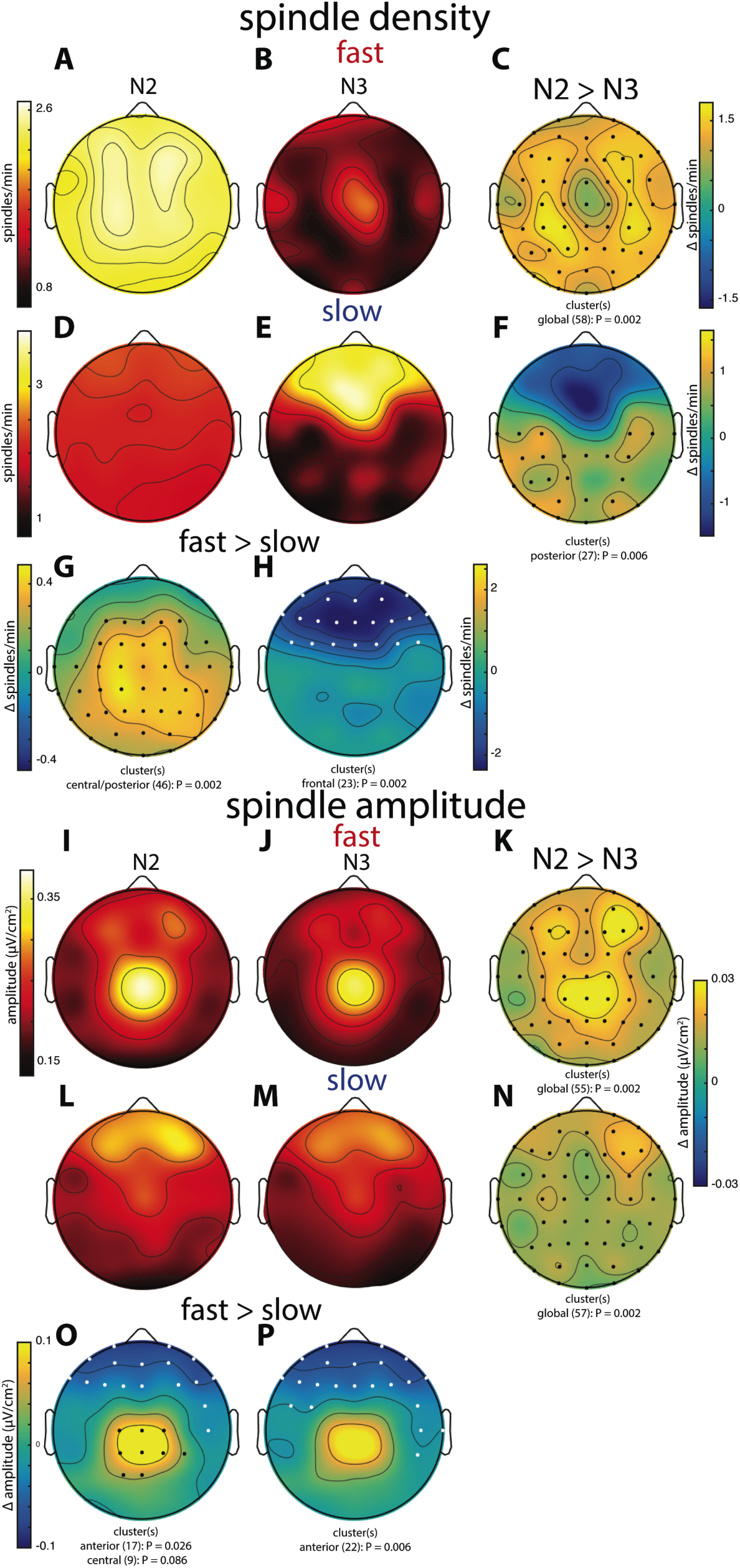
Group topographies of fast and slow spindle density (A-H) and spindle amplitude (I-P) during N2 and N3. (A, I) N2 fast spindles. (B, J) N3 fast spindles. (C, K) N2 > N3 fast spindles difference. (D, L) N2 slow spindles. (E, M) N3 slow spindles. (F, N) N2 > N3 slow spindles difference. (G,O) N2 fast > slow difference. (H,P) N3 fast > slow difference. Significant electrodes from paired tests with cluster correction indicated on difference maps as black (positive) and white (negative) dots. Cluster size(s) (number of electrodes) and P value(s) indicated below each difference map. Note color scales are shared for (A, B), (D, E), (I, J, L, M), (K, N), and (O, P).

Combined, these group-level topographies provide some insights regarding which aspects of spindle activity contribute to observed sigma power. In particular, the N2 fast sigma power peak over centro-parietal regions (Fig. 3) appears to be predominantly driven by enhanced spindle amplitude in this region, but not by spindle density, which was quite uniform across the scalp. In contrast, the bilateral fronto-central profile of N2 slow sigma seems inexplicable in terms of either spindle density or amplitude. Interestingly, both density and amplitude of spindles appear to contribute to the patterns of both fast and slow sigma power in N3.

To examine these group-level observations in more detail, we turned to spatial maps of spindle properties from individual subjects. Similar to sigma power, individual subjects’ spindle density and spindle amplitude profiles were quite variable, and these, too, were quite stable across nights (not shown). We then compared topographical patterns of all three metrics to determine how well they correspond with one another within individuals. For illustration, Fig. 7 shows topographies from one subject for all metrics, for both fast N2 spindles (Fig. 7A) and slow N3 spindles (Fig. 7B). In line with the group maps, this subject’s N2 fast sigma power distribution appears to be more strongly driven by spindle amplitude, and to a lesser extent, if at all, by spindle density (Fig. 7A). In contrast, the slow N3 power topography appears similarly related to spindle amplitude and spindle density profiles (Fig. 7B). These observations suggest that amplitude and prevalence of spindles may contribute to observed sigma power in different ways depending on spindle class and sleep stage.

**Figure 7.**
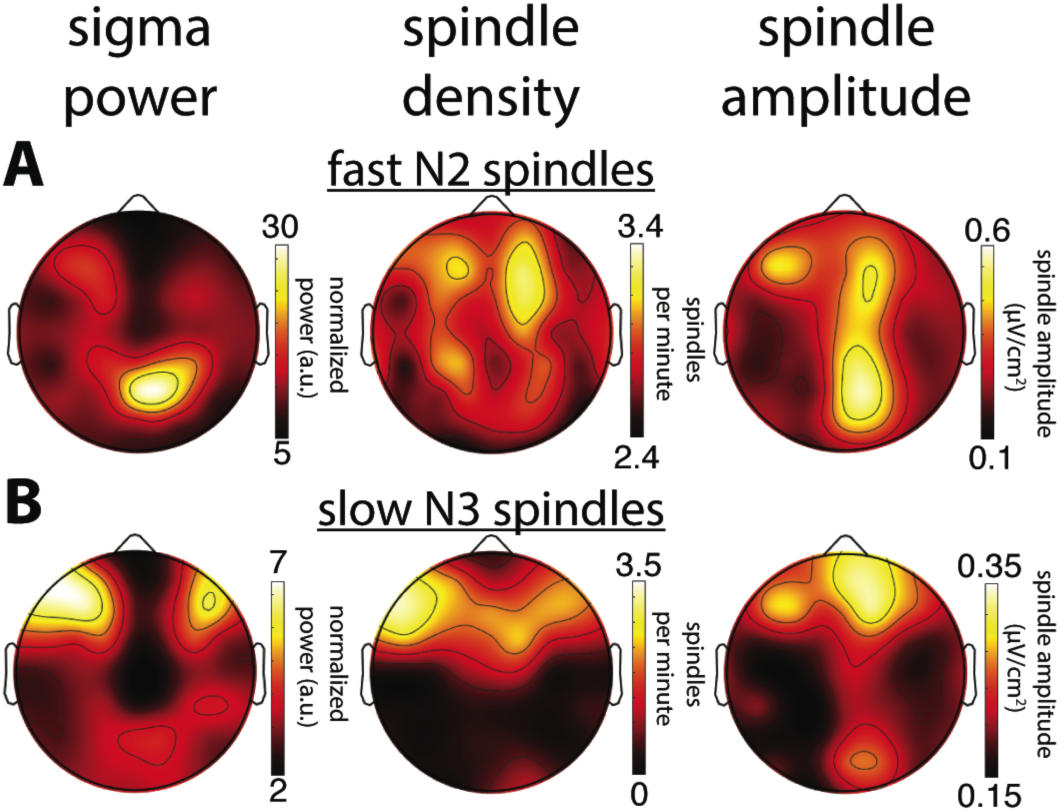
Comparison of topographical maps based on different spindle activity measures for a single subject (S1). (A) Topographies of sigma power, spindle density and spindle amplitude for fast spindles during N2. (B) Topographies for slow spindles during N3. While this subject’s topographies based on different measures show a reasonable correspondence for slow N3 spindles, spatial agreement is quite poor for fast N2 spindles.

To quantify these comparisons, we directly compared topographical patterns between each pair of these three metrics within individuals. As for Table 1, we report correlation coefficients as a measure of topographical similarity, along with classifier recognition performance to indicate whether within-subject similarity exceeds similarity scores seen between subjects.

Detailed values and statistics can be found in Table 2. Overall, within-subject spatial maps based on different metrics showed moderate correspondence, although results depended importantly on sleep stage, spindle class, and which metrics were being compared. Confirming the observations from Fig. 3, Fig. 5 and Fig. 7, fast sigma power topographies were more closely related to spindle amplitude than spindle density for both N2 (t(25)=3.3, P=0.003) and N3 (t(17)=2.9, P=0.01). In contrast, slow sigma power topographies were equally poorly related to spindle amplitude and density during N2 (t(23)=-0.1, P=0.95), but more similar to spindle density than spindle amplitude topographies during N3 (t(18)=-2.0, P=0.06). In terms of subject discriminability, cross-metric subject recognition rates typically did not exceed 40%, except for N3 sigma power vs. spindle density topographies. To place these findings in context, we achieved higher recognition performance for the majority of crosssleep stage and cross-night comparisons based on the same metric (i.e., sigma power), as presented in the previous section (Table 1, Fig. 4 and Fig. 5).

**Table 2.**
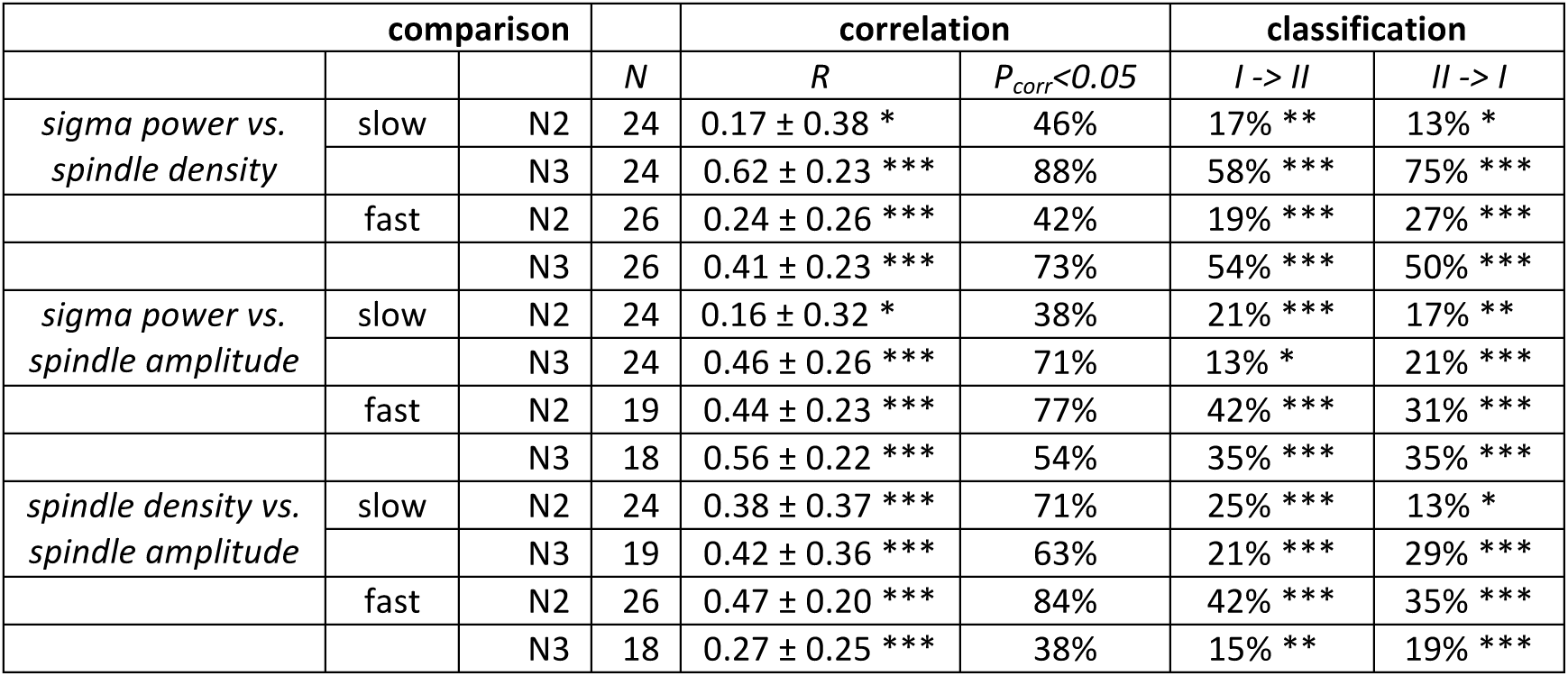
Within-subject similarity of topographies based on different measures of spindle activity. R: Pearson correlation coefficient; P_corr_<0.05: percentage of subjects with False Discovery Rate-corrected P values < 0.05; I-> II and II->I: classification direction. Significance levels (correlation: one-sample t-test vs. zero; classification: binomial test) indicated by *:< 0.05; **:< 0.01; ***:< 0.001.

The spindle metric comparisons lead to two conclusions. First, spindle amplitude and spindle density contribute differently to sigma power, depending on sleep stage and spindle class under consideration. Second, topographical maps based on different metrics are generally quite distinct even for the same individual. This somewhat unexpected finding suggests that the choice of spindle metric could have important downstream ramifications regarding topographical group maps, statistics, and interpretations.

In sum, the analyses in this section again show that group-level maps obscure substantial individual topographical variability of spindle expression. Beyond that, examination of only group topographies would likely lead to the conclusion that different spindle metrics capture very similar aspects of spindle activity. However, more detailed analyses reveal important topographical differences between metrics on the single-subject level, further indicating that group maps mask important features of the oscillatory phenomena we are attempting to understand.

## Discussion

The current study characterized the large between-subject variability of spindle frequencies and topographies, and assessed the correspondence of spatial spindle expression between slow and fast spindle classes, sleep stages, nights, and several metrics of spindle activity. Employing a novel spatial filtering approach to isolate subject-specific frequencies of slow and fast spindle activity, we replicated topographical properties of spindle expression at the group-level^29,30^, strongly suggesting that we successfully isolated the oscillatory phenomena of interest. By then taking a more detailed look at topographical aspects of these oscillatory dynamics, and, furthermore, by extending the subject-specific approach from the spectral to the spatial dimension, we made several novel observations about the organization of NREM spindle dynamics.

### Individual differences in NREM spectra and sigma frequencies

We observed marked individual differences in the shape of NREM power spectra (Fig. 1). Analyses across nights confirmed several previous studies demonstrating the existence of robust spectral power fingerprints during sleep^34,61^. Indeed, these profiles have been shown to be the highly heritable^59^ and remain stable throughout development^62^. In sharp contrast, we observed that N2 and N3 spectral profiles for the same individual could show considerable differences within the same night. While spectral differences between N2 and N3 are inevitable given that they are defined based on the number and amplitude of slow waves, it is perhaps surprising that N2 and N3 spectral profiles are so different.

Indeed, one of the most prominent differences between N2 and N3 spectra appeared in the sigma range, and these differences, too, were highly stable across nights (Fig. 1AC). While fast sigma peaks were more pronounced in N2 than N3 across most individuals, slow sigma peaks did not show consistent amplitude differences between N2 and N3. Yet, both individual sigma peak frequencies and individual topographical patterns of sigma power were highly stable across N2 and N3, suggesting that while the broader oscillatory context may change markedly from light to deep sleep, underlying thalamic spindle generators and their influence on neocortex remain relatively fixed. Thus, the extent to which N2 and N3 are seen to be similar depends on which physiological aspects one considers.

The most striking observation from the spectral profiles, however, was the large between-subject variability of sigma range frequencies, and the difficulty this poses for defining slow and fast sigma ranges that can provide adequate group-level separation of fast and slow spindles. Component-based slow and fast sigma frequencies ranged from 9.3 to 12.0 Hz for slow spindles and from 12.5 to 15.4 Hz for fast spindles, similar to what has been described previously^34,35^. For this reason, it seems any attempt to define spindle activity in terms of fixed spectral criteria is destined to fail in at least some instances. Thus, rather than arguing in favor of specific spectral definitions for slow and fast spindle classes, we believe this variability provides motivation to identify slow and fast sigma peaks in a subject-specific manner^35,39^. However, we demonstrated that both channel-averaged spectra and spectra derived from single frontal or parietal channels are inadequate in this regard, and in many instances do not lead to reliable estimates of subject-specific sigma frequencies.

### Separating slow and fast sigma peaks via spatial filters

To improve upon channel-based approaches, we employed a novel data-driven technique to identify individualized slow and fast sigma peaks. By making use of the fact that multi-channel EEG data has a spectral structure that is typically correlated between nearby channels, we constructed spatial filters that maximally enhance spectral content in one frequency range at the expense of another. This technique is ideally suited to deal with unknown individual differences in topographical spindle expression, which we demonstrated to be substantial. The computational generalized eigendecomposition (GED) backbone we employed has been used previously in various electrophysiological contexts, typically to identify GED components that maximize spectral power in one frequency band relative to broadband activity^52,54,55^. We extend this approach by directly contrasting two closely spaced frequency bands. Importantly, this technique can only provide meaningful separation of spectral components when the oscillatory bands of interest have different spatial profiles. While we used this technique only to determine slow and fast sigma frequencies (and not for topographical analyses *per se*), successful separation of slow and fast sigma peaks in virtually every subject implies that these oscillatory components are sufficiently spatially distinct.

When we compared component-derived spectral peaks to both channel-averaged and single-channel (frontal and parietal) spectra, we found a clear advantage for the spatial filter approach. Compared to peaks isolated from individual channels, the component-based approach allowed detection of slow sigma peaks in approximately three-quarters of the sample that would have otherwise been lost for downstream analyses. Nonetheless, it is important to note that channel-and component-based sigma peaks were significantly correlated across individuals (Supplementary Text), reflecting the obvious dependence of GED-derived spindle peaks on individual channel information.

Several other approaches have been proposed to identify subject-specific sigma ranges. For example, the individual adjustment method^35,39^ averages the second derivative of each channel’s power spectrum across channels to determine individualized slow and fast spindle ranges, while other research groups average spectra separately across anterior and posterior channels to identify slow and fast peaks^26^. While we did not directly compare our technique to these approaches, we see important benefits to the GED-based spatial filtering approach. First, because it is data-driven, it makes no *a priori* assumptions regarding which channels exhibit slow or fast spindle activity. Indeed, individual topographical maps exhibited substantial variability in local spindle expression, such that anterior vs. posterior channel selection may not always successfully resolve distinct peaks (see, for example, Fig. 4D). Second, our approach consistently returned highly similar slow sigma frequencies for N2 and N3. This contrasts with observations of less well-matched slow sigma peaks in N2 and N3 when using spectra averaged across groups of channels^26^, or single-channel spectra (this study). Likely, this poor correspondence is due to fast sigma peaks overshadowing slow sigma peaks during N2, an issue that the GED approach inherently tries to avoid by increasing spectral power in one sigma band relative to the other. Taken together, we suggest our method is a more sensitive technique to robustly identify individuals’ fast and especially slow spindle frequencies than those previously described. Based on this validation of the GED-based approach, we analyzed the spatial characteristics of sleep spindles.

### Sleep spindle topographies

We analyzed topographical patterns of spindle expression both at the group and individual level. Group-wise, we observed typical differential topographies of slow and fast spindles, suggesting we succeeded in separating the two spindle classes. Canonical slow-frontal vs. fast-central topographies were observed in N3 for all spindle metrics (Fig. 3 and Fig. 6). In N2, we also observed a central expression of fast spindle activity for sigma power and spindle amplitude, but a less typical fronto-central topography for slow sigma power, and a relatively uniform distribution of both slow and fast spindle densities. Direct statistical comparisons of slow and fast spindles supported distinct topographies for most sleep stage/spindle class/spindle metric combinations. We also observed stronger fast spindle activity during N2 than N3, regardless of metric, consistent with earlier observations^30^. In contrast, frontal slow sigma power and slow spindle density were greater during N3 than N2, consistent with more pronounced slow sigma spectral peaks in N3 than N2. Here, it should be noted that N2 slow spindles have not been examined in detail previously, likely because channel-based spectra did not unambiguously indicate their presence during N2. Thus, the increased sensitivity afforded by targeting subject-specific spindle frequencies uncovers subtle differences in sigma power expression during N2 and N3 sleep that might otherwise be overlooked.

When we examined in detail individual topographical profiles of spindle expression, we observed substantial individual differences (Fig. 4). Importantly, individual topographical variability was stable both across sleep stages and nights (Fig. 5), analogous to topographical EEG stability during wakefulness^31^, indicating these patterns constitute robust traits. Together, these findings underscore that group averages reflect a mixture of quite idiosyncratic spatial spindle patterns.

What could be the cause of such marked individual topographical variability? First, it is possible that spatial variability is due to individual differences in cortical folding patterns, affecting how intracortical spindle signals summate and propagate to the scalp. Second, different individuals might have different anatomical or functional thalamocortical wiring profiles, resulting in different spatial profiles of spindle expression. Indeed, between-subject variability in white matter tracts, including fibers intrinsic to and surrounding the thalamus, is known to be related to spindle power and density^63^. More generally, there is substantial evidence for the existence of localized cortical and subcortical spindles^4–6,64^ and the potential role of localized spindles in processing specific memories^18,65^–68. Thus, an intriguing possibility is that individual differences in topographical spindle expression have functional implications and underlie individual differences in cognitive processing.

Regardless of the question of function, topographical heterogeneity may be a source of substantial noise in research designs choosing a specific channel for spindle analyses. For example, rank-ordering subjects according to sigma power on channel Cz will, in all likelihood, yield a different order from one based on each individual’s channel of maximum sigma power. As a result, differences between experimental or clinical groups could stem from these groups having different cortical regions with maximal spindle activity, either systematically or by chance. Similarly, experimental studies testing topographical hypotheses might benefit from taking spatial variability into consideration by recording a baseline night prior to experimental manipulation.

### Not all spindle metrics are alike

Beyond differences in spectral definition, studies of spindles report a host of different metrics intended to capture (specific aspects of) underlying spindle activity. Classical measures of sigma power are reported along with various properties derived from individually detected spindles (e.g., spindle density, amplitude, peak frequency, power, duration, integral, etc.). While it is well understood that different metrics are, in a trivial sense, distinct, they often appear to be lumped together when relating findings from different studies.

When we compared individuals’ spatial topographies between different spindle activity metrics, we made several observations. First, spindle amplitude and spindle density were found to contribute differently to sigma power, depending on sleep stage and spindle class under consideration. Whereas fast sigma power was more strongly driven by spindle amplitude than spindle density across sleep stages, the opposite was found for slow sigma during N3. Second, while individuals’ topographical patterns of sigma power, spindle density, and spindle amplitude showed above-chance levels of correspondence, cross-metric subject recognition was rather low. In general, neither spindle density nor spindle amplitude appeared greatest where sigma power was highest. Similarly, sites where spindles were more plentiful did not necessarily have larger spindles.

How do these various metrics relate? Sigma band power, as a relatively crude measure of spindle activity, likely includes noise unrelated to physiological events of interest, but also captures meaningful activity of low amplitude and short duration spindles that do not register with specialized spindle detection algorithms. Conversely, spindle detection methods are forced to make ultimately arbitrary decisions regarding amplitude and duration criteria, potentially leading to nondetection of relevant spindle activity (see^69,70^ for the difficulties associated with validating automated spindle detectors against human scorers). Anecdotally, we often observed in the raw EEG trace brief, but prominent, increases in sigma amplitude during slow wave up states that were abruptly suppressed by subsequent down states. In many cases, however, these sigma events were shorter than the 400 ms criterion employed in our current algorithm and were not counted as spindles. In the absence of ground truth, we have no clear preference for taking a sigma power or spindle detection approach, nor do we advocate specific spindle detection criteria or particular spindle metrics. However, as our findings show, it is important to realize that different metrics capture different components of spindle activity and cannot be assumed to correspond to one another in a straightforward fashion.

### On the existence of slow and fast spindles

While the evidence for distinct slow and fast spindles is surprisingly equivocal in the animal literature^64,71,72^, our findings are well in line with the current notion of two distinct types of human spindles^8,24–28^. Indeed, our group-level topographical maps agree with previous reports^29,30^. More fundamentally, we were able to identify separate, and typically quite narrow, slow and fast spectral peaks using the GED spatial filter approach for every individual (although we did exclude three individuals with inconsistent N2 and N3 slow peaks). It is worth emphasizing that the GED spatial filter approach does not artifactually introduce peaks, as there were still some instances where we could not identify sigma peaks from component spectra. Moreover, the clear congruence of component peak locations between nights and sleep stages argues against such methodological explanations.

Previous studies described the existence of an anterior-posterior gradient of spindle frequency^5,73^ but it is unclear how the distinct spectral peaks found here could have emerged from a more continuous distribution of sleep spindle frequencies. Instead, such findings may be wholly explained in the framework of a bimodal division of slow and fast spindles. As others^27,29,30^ and we have shown, both slow and fast spindles occur widely across the brain, but with more anterior or posterior biases. Thus, averaging over many spindles (or, regarding sigma power, many spectra derived from short data segments) can give rise to a spatial frequency gradient by sampling different proportions of slow and fast spindles at different cortical sites.

Similarly, findings of varying spindle frequencies throughout the night or within sleep cycles^30,74,75^, may be explained by shifting contributions of slow and fast spindle activity as opposed to the occurrence of progressively slower or faster spindles. Indeed, while GED-based sigma frequencies were stable from N2 to N3 (Fig. 2B), channel-based frequencies were much more variable between sleep stages (Fig. 2F), indicating that the same underlying combination of slow and fast sigma generators can manifest in the power spectrum as peaks with different locations. In sum, we believe the current evidence, including our findings, argues in favor of two fundamentally distinct spindle classes in humans.

### Practical recommendations

We offer several suggestions regarding slow and fast spindle separation. First, we believe the field would benefit from targeting subject-specific frequencies. While it was outside the scope of the current work to directly compare individualized spectral bands to a fixed frequency approach^35^, we hope the variability showcased in Fig. 1 makes the point. Individuals with highest sigma power levels based on fixed spectral criteria are not necessarily the ones with greatest power when using individually defined sigma frequencies. Fixed frequency criteria could therefore have undesired effects, e.g., when assessing the link between spindle activity and memory. As another concern, even small changes in sigma range settings can have comparatively large effects on an individual’s topography, as "true" slow and fast topographies are differentially enhanced, attenuated, and mixed. While we hope our spatial filtering approach will prove useful in this regard (see Supplementary Materials for Matlab code implementing the GED-based technique), the precise method employed is less important than adopting an individualized approach in the first place.

Second, when subject-specific frequency selection is not feasible, we suggest fixed criteria be based on visual inspection of power spectra. In our sample, approximate slow and fast spindle ranges of 9-12.5 Hz and 12.5-16 Hz would likely have worked reasonably well. Moreover, based on visual inspection of the distribution of peak frequencies in a sample of 161 individuals^35^ (their Fig. 2) these ranges appear to be a sensible choice for young, healthy individuals. However, we caution against blindly following this suggestion, as slow and fast sigma peak distributions in other samples might overlap more^35^, or might be clustered in different spectral bands in different age ranges or clinical groups^76^. Still, we emphasize adopting fixed criteria constitutes an unnecessary reduction in statistical power and may lead to spurious findings in smaller samples.

Finally, we recommend careful examination of spindle detection criteria used in the literature. Study conclusions regarding the specific involvement of slow or fast spindles that appear at odds may no longer be when one considers in detail the spectral criteria employed. Specifically, we suggest that studies placing lower bounds for slow spindles at 11 or 12 Hz may have missed their slow spindle target in some subjects, and/or confused fast for slow spindles by artificially separating the fast sigma range into two. Rather than criticizing these studies, however, we simply wish to alert researchers and clinicians that seemingly minor methodological details could have major impact on the interpretation of results.

### Conclusion

Ever since their first description^77^, sleep spindles have been a topic of considerable interest^10,11,78–80^. This interest has only increased in recent years with accumulating evidence of the role of sleep spindles in memory consolidation^13–16^, and of their dysfunction in several neuropsychiatric conditions^32, 40, 81–83^. Despite decades of progress, the functional role of sleep spindles in general, and of slow and fast spindles in particular, is still unclear. While some recent evidence suggests fast spindles are more strongly implicated in cognitive processes^17–22^, only few studies finding such links target the purported brain rhythms in an individualized manner^26^. But if slow and fast spindles do indeed turn out to serve different functions, to serve the same function differently, or to be differentially affected in neuropsychiatric disorders, it becomes critical that these spindle types be adequately separated. We suggest that proper understanding of sleep spindle dynamics relies on acknowledging the distinct properties of slow and fast spindles in light and deep sleep, and on addressing individual variability as much as accounting for robust group-level phenomena. We hope this approach will prove a useful foundation for future investigations into the physiological properties of sleep spindles and their functional role in both healthy and clinical states.

## Acknowledgements

This work was supported by grants from The Netherlands Organization for Scientific Research (NWO) to RC (446-14-009); National Institutes of Health to ACS (F32-NS093901), RS (MH048832), DSM (K24MH099421), RS and DSM (MH092638); The Harvard Clinical and Translational Science Center (TR001102); and Stanley Research Center. We thank Bengi Baran, Cameron Callahan, David Correll, Charmaine Demanuele, Rachel Fowler, Elaine Parr, Ben Seicol and Tessa Vuper for their data collection and preprocessing efforts.

